# An Evolutionary Novelty in TRPV1 Functional Regulation: Characterization of a Dominant-Negative Isoform Exclusive to Catarrhine Primates

**DOI:** 10.1101/2025.01.24.634766

**Authors:** Sofía Cancino, Deny Cabeza-Bratesco, Ella Matamala, Julia M. Jork, Kattina Zavala, Cristian Montecinos, Sebastián Bustamante-Villarreal, Rodrigo Maldonado, Gonzalo Riadi, Sebastian E. Brauchi, Juan C. Opazo

## Abstract

TRPV1, a member of the transient receptor potential (TRP) family, is a non-selective cation channel primarily known for its role in pain perception, inflammation, and thermosensation. In mammals, it responds to noxious heat (>43°C) and chemical stimuli such as capsaicin and protons. It is widely expressed in sensory neurons, notably in the dorsal root and trigeminal ganglia. However, it is also found in some non-neuronal tissues, like the skin and bladder. The human canonical variant of TRPV1 renders a protein with 839 residues. Different splice variants have been described, and some display a dominant negative effect, partially or totally inhibiting the activity of the canonical counterpart. Here, we characterize a splice variant that encodes for a channel of 850 amino acids (TRPV1_850_). This variant is an evolutionary novelty of catarrhine (Old World monkeys and apes) primates, incorporating an exon of 33 bp long. Both imaging of membrane expression and electrophysiological recordings suggest that TRPV1_850_ alone does not reach the plasma membrane. However, in cells co-expressing the canonical and TRPV1_850_ variants, the latter would act as a dominant negative, preventing the canonical variant from reaching the plasma membrane and rendering smaller macroscopic currents in response to capsaicin. Thus, this new isoform of the TRPV1 ion channel represents a novel form of functional regulation only present in apes and Old World monkeys.

**Significance:** We identified a novel isoform of the TRPV1 ion channel, a protein essential for detecting pain and heat stimuli unique to apes and Old World monkeys. This isoform contains an additional exonic sequence encoding eleven amino acids, which originated in the common ancestor of apes and Old World monkeys. This additional exonic sequence disrupts a specific domain in the N-terminal region of the protein, resulting in functional differences. Unlike the canonical TRPV1 isoform, this variant cannot reach the plasma membrane independently. Instead, it exerts a dominant-negative effect by interfering with the canonical TRPV1 protein, reducing its ability to localize to the cell surface and diminishing its responsiveness to capsaicin, a well-known TRPV1 activator.

## Introduction

Proteins play a central role in life. With the advancement of sequencing technology, it is possible to know precisely how many protein-coding genes are present in our genomes. According to ENSEMBL v.113, the primary assembly of the human genome possesses almost 20 thousand protein-coding genes (Harrison et al. 2024). However, the repertoire of proteins that our genomes can produce is more than this number. This is possible due to the alternative splicing process, which selectively includes or excludes exons during the RNA processing (Berget et al. 1977; Chow et al. 1977). According to the most recent estimations, each human protein-coding gene can produce, on average, almost 20 different isoforms (Harrison et al. 2024). There are examples showing that different splice variants can be expressed at different developmental times, subcellular localizations, and/or in different tissues (Ozaita et al. 2002; Senatore et al. 2014; Liang et al. 2021; Guan et al. 2022), highlighting the power of the alternative splicing process in generating functional diversity. Additionally, clade-specific emergence of new exons further increases the functional repertoire from a single gene in a specific group of species (Kondrashov and Koonin 2001; Alekseyenko et al. 2007; Merkin et al. 2015).

TRPV1 (transient receptor potential vanilloid 1) is a non-selective cation channel primarily activated by heat (>43°C), capsaicin, and acidic pH (Rosenbaum and Islas 2023). It plays a crucial role in nociception and thermosensation (Caterina et al. 1997). Also known as VR1, Osm-9-like-TRP, and capsaicin receptor, TRPV1 is also found in non-neuronal tissues, including skin, bladder, and vasculature (Szallasi et al. 2007). Its subcellular trafficking and plasma membrane expression are tightly regulated, with mechanisms involving endocytic recycling, phosphorylation, and interaction with scaffolding proteins (Ferrandiz-Huertas et al. 2014; Rivera et al. 2024). These processes affect its localization at the plasma membrane, impacting receptor availability and sensitivity to stimuli. It has been shown that the strong tachyphylaxis observed in TRPV1, reflected by an acute decrease in response to capsaicin administration, can partly be explained by trafficking control of the channel (Tian et al. 2019).

Several splice variants of the TRPV1 ion channel have been described, including TRPV1b and TRPV1α (Sharif Naeini et al. 2006; Vázquez and Valverde 2006; Schumacher and Eilers 2010). Common human splice variants result in a truncated protein with altered functional properties and plasma membrane expression (Schumacher et al. 2000). For example, TRPV1b forms functional ion channels that are activated only by temperature, not by capsaicin or protons, acting as a dominant-negative, reducing canonical TRPV1 activity when co-expressed, likely by forming heteromers that diminish channel response to activation (Lu et al. 2005). Other variants may exhibit distinct subcellular localizations and activity levels, influencing the overall functional repertoire of TRPV1 in tissues where they co-occur (Wang et al. 2004).

In this work, we discovered a 33 bp exon in the gene encoding the TRPV1 ion channel that is only present in catarrhine primates. This exon disrupts the sixth ankyrin domain, and its incorporation originates a splice variant of 850 amino acids (TRPV1_850_). We find this isoform does not traffic to the plasma membrane, and when coexpressed with the canonical isoform, it effectively retains TRPV1_839_ intracellularly, displaying a smaller current density when activated by capsaicin without affecting the open probability of the channel. Hence, from a functional perspective, the isoform of 850 amino acids modulates the cellular response to capsaicin. According to our preliminary assessment of tissue expression, this isoform has a narrow gene expression pattern, so the dominant negative effect should be relevant only in a defined subset of tissues. Thus, the exon gain in the last common ancestor of apes and Old World monkeys represents an evolutionary innovation related to the negative regulation of the TRPV1 ion channel.

## Results and Discussion

### Coding exon gain in the last common ancestor of apes and Old World monkeys

Our manual annotation of the TRPV1 channel found an exonic sequence in the sixth position in apes and Old World monkeys (Fig. 1). This exon has a length of 33 nucleotides, coding for an 11 amino acid sequence (Fig. 1). We also identify this sequence in New World monkeys, however in all examined species the sequence has a single nucleotide deletion (Fig. 1). This deletion was found in the same position in all examined New World monkeys, suggesting that it was present in the last common ancestor of the group (Fig. 1). This single nucleotide deletion produces a shift in the reading frame originating several stop codons. The first premature stop codon was identified at position 410 of the protein (Fig. 2) before the first transmembrane domain annotated for the human TRPV1 ion channel, which is located between position 445 and 465 of the splice variant of 850 amino acids (TRPV1_850_). Thus, our results suggest this sequence does not form an encoding exon in New World monkeys. In the case of tarsiers and strepsirrhines, the third codon of the sequence is a stop codon (Fig. 1). Therefore, in the case of incorporating the sixth exon, the cellular machinery of tarsiers and strepsirrhines is hypothesized to produce an even shorter protein in comparison to New World monkeys also with no transmembrane domains. Accordingly, our results suggest that this sequence is also not an encoding one for the TRPV1 ion channel in tarsiers and strepsirrhines. In the case of representative species of the mammalian orders that are closely related to primates, Scandentia (treeshrews) and Dermoptera (colugos) (Mason et al. 2016), they also have a stop codon in the same position as tarsiers and strepsirrhines (Fig. 1). A similar situation was observed in the case of the house mouse (Fig. 1).

**Figure 1.**
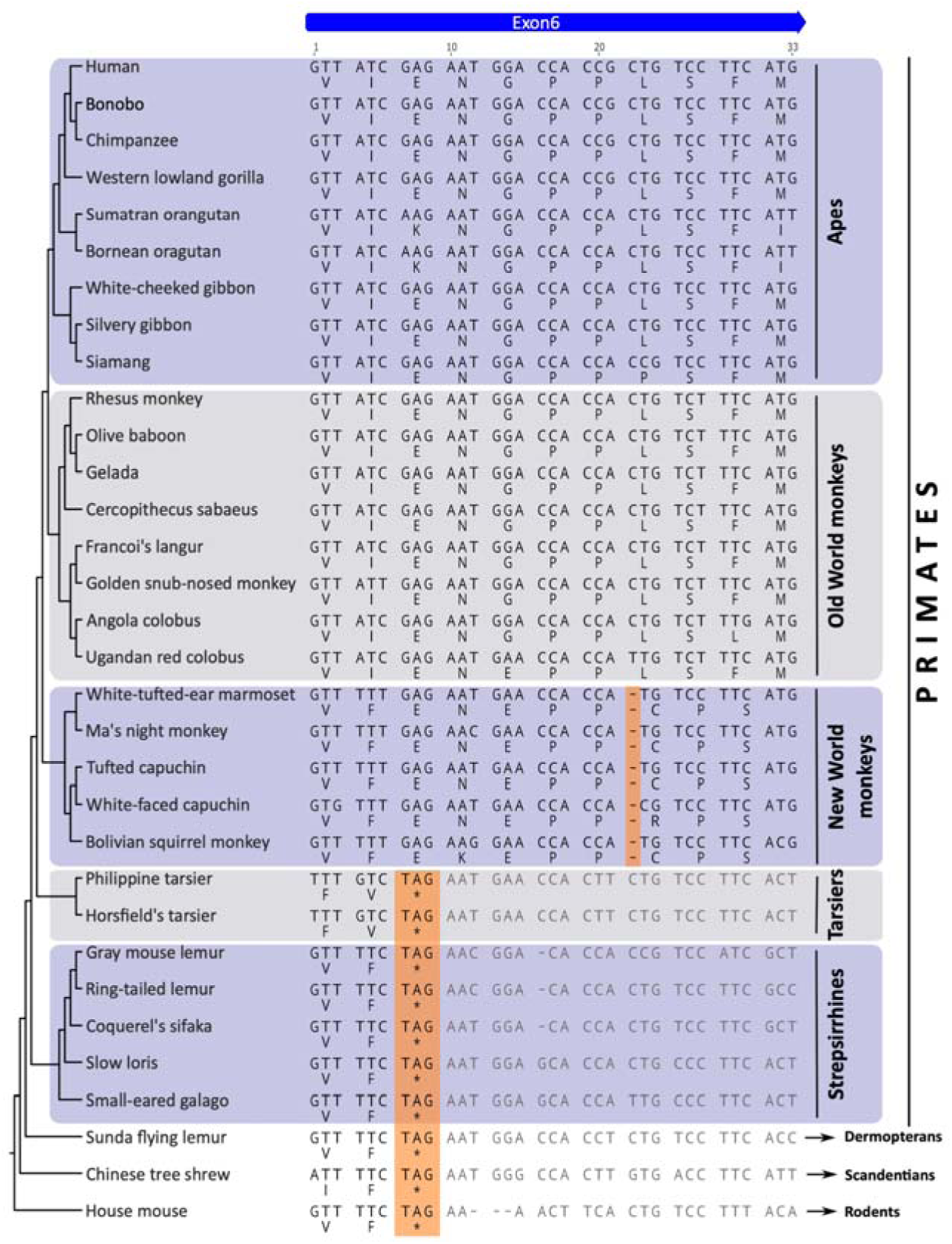
Nucleotide/amino acid alignment of the sequence of the sixth exon of the TRPV1 ion channel in apes and Old World monkeys with the corresponding sequence in other primates, dermopterans, scandentians, and rodents. The orange shading highlights the single nucleotide deletion in New World monkeys and the stop codon in tarsiers, strepsirrhines, dermopterans, scandentians, and rodents. Nucleotide sequences were aligned using the program MAFFT v7 (Katoh et al. 2019). Phylogenetic relationships among the species were obtained from the literature (Perelman et al. 2011; Mason et al. 2016; Shao et al. 2023).

**Figure 2.**
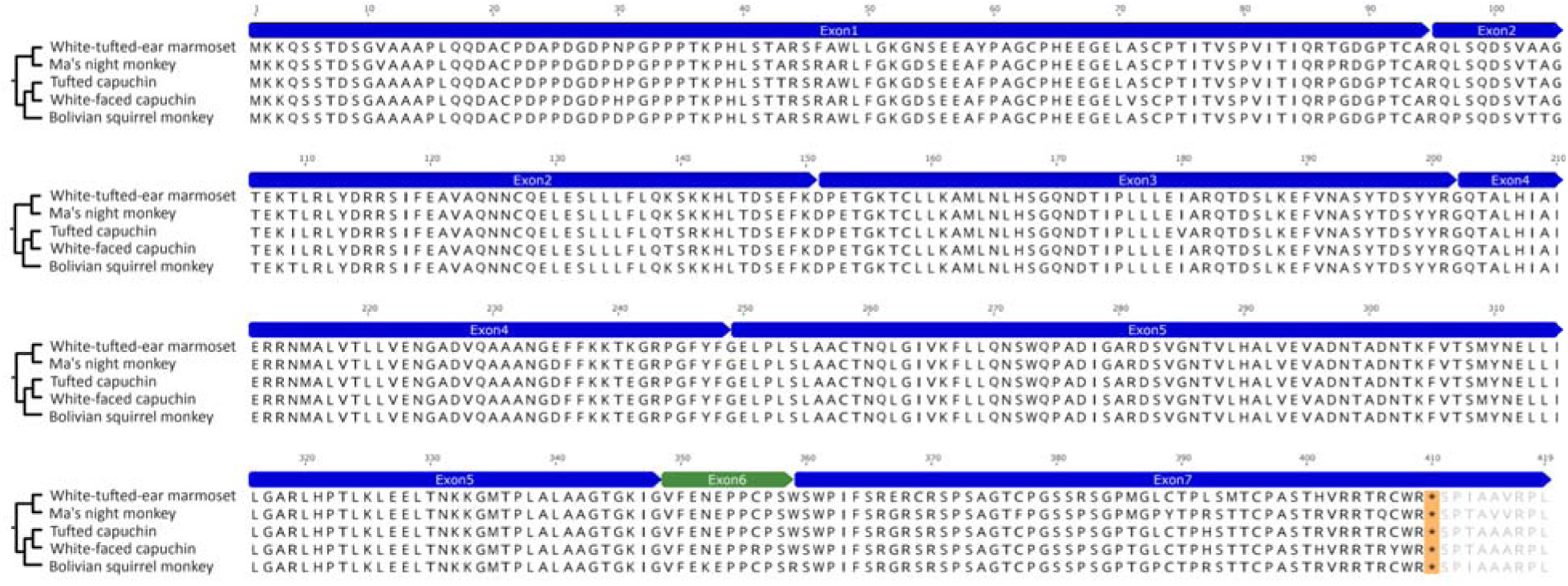
Amino acid alignment of the TRPV1 ion channel in New World monkeys incorporating the sequence of the sixth exon (green). The orange shading highlights the first premature stop codon at position 410, which is before the first transmembrane domain located between position 445 and 465 of the splice variant of 850 amino acids (TRPV1_850_), according to the UniProt database (UniProt Consortium 2023). Amino acid sequences were aligned using the program MAFFT v7 (Katoh et al. 2019). Phylogenetic relationships among the species were obtained from the literature (Perelman et al. 2011; Mason et al. 2016; Shao et al. 2023).

The evolutionary origin of new exons provides an opportunity to have a more diverse repertoire of protein isoforms from a single gene through alternative splicing (Kondrashov and Koonin 2001; Alekseyenko et al. 2007; Merkin et al. 2015). This process is a fundamental mechanism in evolution, increasing the functional complexity of genomes. It permits organisms to evolve biological traits without increasing the number of genes, allowing evolution to exploit the potential of existing genetic information (Singh and Ahi 2022; Verta and Jacobs 2022). According to the Ensembl database v.113 (Harrison et al. 2024), the number of transcripts for the human TRPV1 ion channel (7 transcripts) is lower in comparison to the average number of transcripts per gene based on the primary assembly of the human genome (19.5 transcripts per gene). This finding could be explained by the fact that large gene families produce fewer splice variants (Kopelman et al. 2005). According to Kopelman et al. (2005), large gene families, i.e., more than ten paralogs, use alternative splicing approximately half as frequently as gene families with one family member. Alternative splicing plays a fundamental role in ion channel physiology. There are several examples showing that ion channel isoforms are expressed at different developmental times, subcellular localizations, and/or in different tissues; also, different ion channel isoforms can have different ion permeability (Ozaita et al. 2002; Senatore et al. 2014; Liang et al. 2021; Guan et al. 2022). The case of the great pond snail (*Lymnaea stagnalis*) is remarkable, as its Ca_V_3 T-type ion channel has isoforms that are expressed in different tissues exhibiting different ion permeability (Ca^+2^ or Na^+^) (Senatore et al. 2014; Guan et al. 2022).

Thus, our results suggest that the sixth exon of the TRPV1 ion channel is an evolutionary innovation of apes and Old World monkeys that originated in the ancestor of the group between 43 and 28.8 million years ago (Kumar et al. 2022).

### The amino acid sequence of the sixth exon of the TRPV1 ion channel disrupts the sixth ankyrin domain

To continue characterizing the sixth exon of the TRPV1 ion channel, which originated in the ancestor of apes and Old World monkeys, our next question was to find out where in the protein structure it is located. After annotating all the domains found in the TRPV1 ion channel according to UniProt (UniProt Consortium 2023), we discovered that the amino acid sequence corresponding to the sixth exon disrupts the sixth ankyrin domain (Fig. 3).

**Figure 3.**
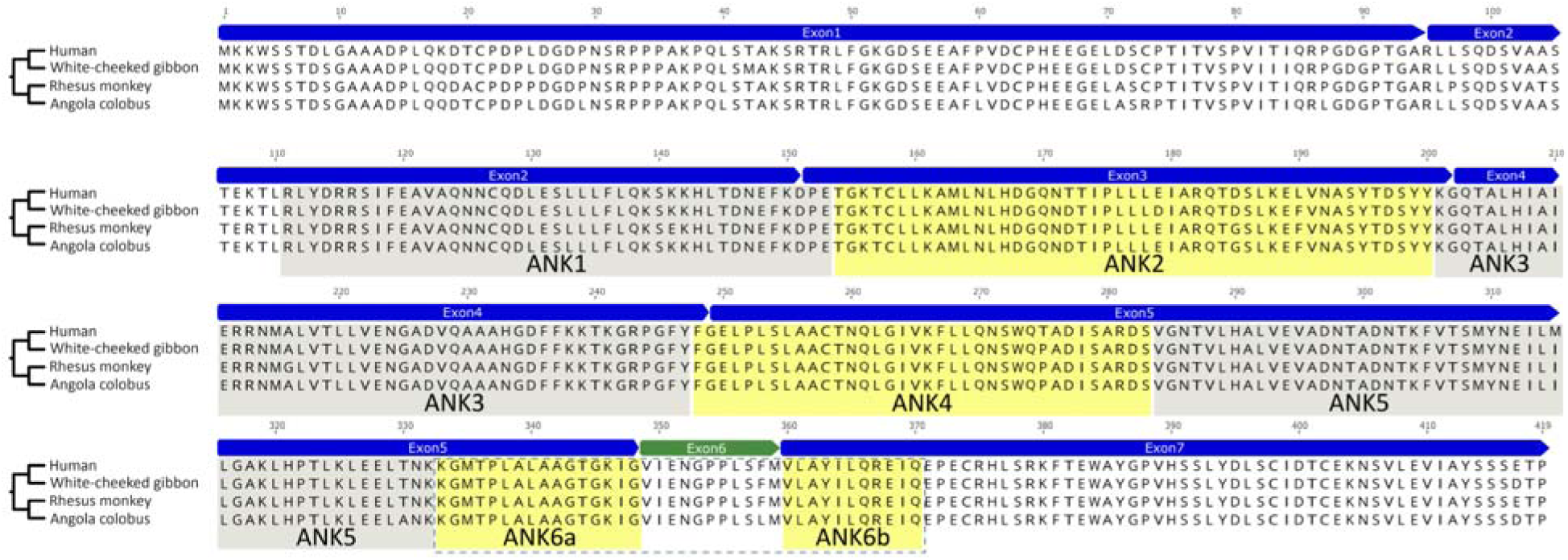
Amino acid alignment of the N-terminal region of the TRPV1 ion channel in representative species of apes (human and white-checked gibbon) and Old World monkeys (rhesus monkey and Angola colobus). The yellow and gray shadings depict the location of the ankyrin domains according to the UniProt database (UniProt Consortium 2023). It is interesting to note the disruption of the sixth ankyrin domain by the 11 amino acid sequence of the sixth exon. Amino acid sequences were aligned using the program MAFFT v7 (Katoh et al. 2019). Phylogenetic relationships among the species were obtained from the literature (Perelman et al. 2011; Mason et al. 2016; Shao et al. 2023).

Ankyrin domains are structural motifs found in proteins in all branches of the tree of life (Gaudet 2008; Chakrabarty and Parekh 2022). Although the number of repeats varies, most proteins possess between two and six repeats (Chakrabarty and Parekh 2022). In the case of the TRPV1 ion channel, six ankyrin domains have been described in the N-terminal portion of the protein with a length that varies between 36 and 49 amino acids (UniProt Consortium 2023). They play a crucial role in the function and regulation of the TRPV1 ion channel. Ankyrin repeats serve as protein-protein interaction sites, facilitating the assembly of signaling complexes and modulating their activity (Mosavi et al. 2004; Gaudet 2008; Brauchi 2023). These domains are also important for the correct channel trafficking to the cell membrane and its sensitivity to diverse intracellular and extracellular factors (Lishko et al. 2007; Ladrón-de-Guevara et al. 2020). For example, it has been demonstrated that a motif of two amino acids in the first ankyrin repeat is essential to determine the activation temperature of the TRPV1 ion channel (Hori et al. 2023). Thus, ankyrin domains contribute to the diverse range of sensory responses mediated by TRPV1, making them a crucial element in understanding the function and regulation of this ion channel.

Finally, after characterizing the presence of this isoform in the genome of apes and Old World monkeys, our next step was to test whether this isoform is transcribed. This seems required, given the challenge of annotating exons of this length (Parada et al. 2021; Yu et al. 2022; Liu et al. 2023). In fact, this exon is annotated in the human genome in Ensembl but not in NCBI. Ensembl annotation comes from ENCODE version 47, which has a file with literature evidence for each transcript in its annotation. However, in the case of this particular transcript, the three supporting works did not confirm the existence of the transcript. Thus, our first approach was to search the NCBI database, where we found a single record expressed in human testis (DQ177333.1). However, this record does not have a corresponding report in the literature. Thus, to confirm the existence of this transcript, we analyzed the RNA-seq data corresponding to 54 tissues for more than 17 thousand human samples available in the Genotype-Tissue Expression database (GTEx) (https://gtexportal.org/home/). This database shows that this isoform has a narrow expression pattern, as we found expression in the testis, cerebellum, and lung (Supplementary Figure S1A-B). To extend our findings, we checked the presence of reads of this splice variant by analyzing 171 selected RNA-seq runs across 27 healthy tissues (Bioproject PRJEB4337) (Fagerberg et al. 2014). After analyzing nearly 3.5TB of sequence data (5 billion reads), we identified reads that met our rigorous criteria (see Methods). These reads were found in five SRA runs from the small intestine (ERR315388; ERR315408), colon (ERR315403), kidney (ERR315443), and testis (ERR315456) biosamples. These results further confirm the existence of this splice variant. Knowing that this isoform is transcribed, our next step was to verify whether the protein exists. To do so, we examined the Peptide Atlas database (https://peptideatlas.org/), a publicly accessible compendium of peptides identified in a large set of tandem mass spectrometry proteomics experiments (Desiere et al. 2006). Our searches show that the protein corresponding to the isoform of 850 amino acids (E7EQ78) is present in different human cells and tissues (Supplementary Figure S1C). Thus, our searches in different databases show that this isoform is transcribed and translated, i.e., exists in nature.

### The splice variant incorporating the sixth exon, TRPV1_850_, does not traffic to the plasma membrane

To determine whether the splice variant that incorporates the sixth exon, TRPV1_850_, displays a similar subcellular distribution as the canonical splice variant, TRPV1_839_, we transiently transfected HEK293T cells with TRPV1_839_ as well as TRPV1_850_, fused with either EGFP or HaloTag. Consistent with previous reports (Senning and Gordon 2015), total internal reflection fluorescence (TIRF) microscopy revealed that TRPV1_839_ fluorescence displays a characteristic signal, spatially distributed in the form of puncta, primarily localized at the plasma membrane (Figure 4A). In contrast, the signal corresponding to TRPV1_850_ was observed as a reticular pattern (Figure 4B) that colocalized well with Sec61b (a marker of the endoplasmic reticulum) (Figure 4C), suggesting that TRPV1_850_ is retained at the endoplasmic reticulum (ER), likely at the cortical ER (Poteser et al. 2016). These results are in agreement with the evidence showing that ankyrin repeats are essential for the correct channel trafficking to the cell membrane (Lishko et al. 2007; Ladrón-de-Guevara et al. 2020), and in particular that the sixth ankyrin domain of the TRPV1 channel is related to that function.

**Figure 4.**
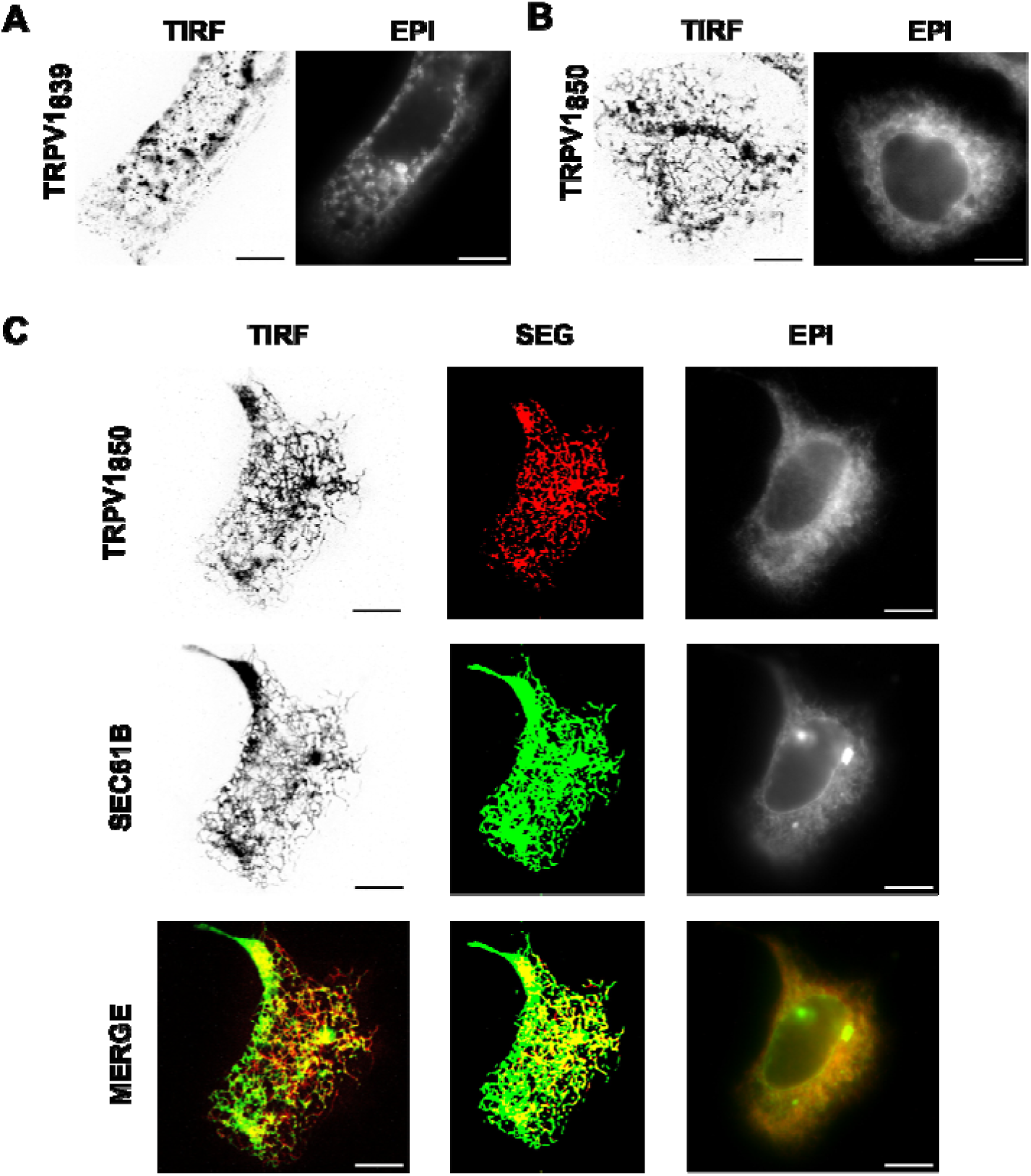
Membrane expression of transfected HEK293 cells with TRPV1_839_ or TRPV1_850_. **A.** Cells transfected with TRPV1_839_ show characteristic puncta (darker spots) at the evanescent field (TIRF) that contrast the diffuse signal observed in regular fluorescence configuration (EPI). **B.** Cells transfected with TRPV1_850_ display a reticular pattern at the TIRF plane and intracellular accumulation (EPI). **C.** Cotransfected cells with TRPV1_850_ and SEC61B (a marker of the endoplasmic reticulum) show high colocalization and similar reticular pattern. The result of the segmentation (SEG) analysis is in the middle.

Interestingly, the cotransfection of both isoforms allows TRPV1_850_ to reach the plasma membrane, while the density of TRPV1_839_ puncta remains unaffected (Figure 5A). To confirm whether the observed puncta was at the membrane and not in a vesicular compartment close to the membrane, we fabricated membrane sheets by sonication (see methods), effectively eliminating false positives by unroofing cells (Toro et al. 2015). Coexpression analysis of the segmented signals from unroofed cells shows that the isoforms likely form heterotetramers where 33.63% (±7.60%) of the TRPV1_839_ signal overlaps with TRPV1_850_, while 53.19% (±10.43%) the of TRPV1_850_ signal overlaps with TRPV1_839_ (Figure 5B). At the same time, we observed a large signal for intracellularly retained TRPV1_839_ (Figure 5B, EPI), suggesting a certain amount of impairment for correct channel trafficking. Overall, our results suggest that the splice variant TRPV1_850_ takes advantage of the natural trafficking of the canonical variant TRPV1_839_ to reach the plasma membrane, and in doing so, it also retains a fraction of TRPV1_839_ at the ER.

**Figure 5.**
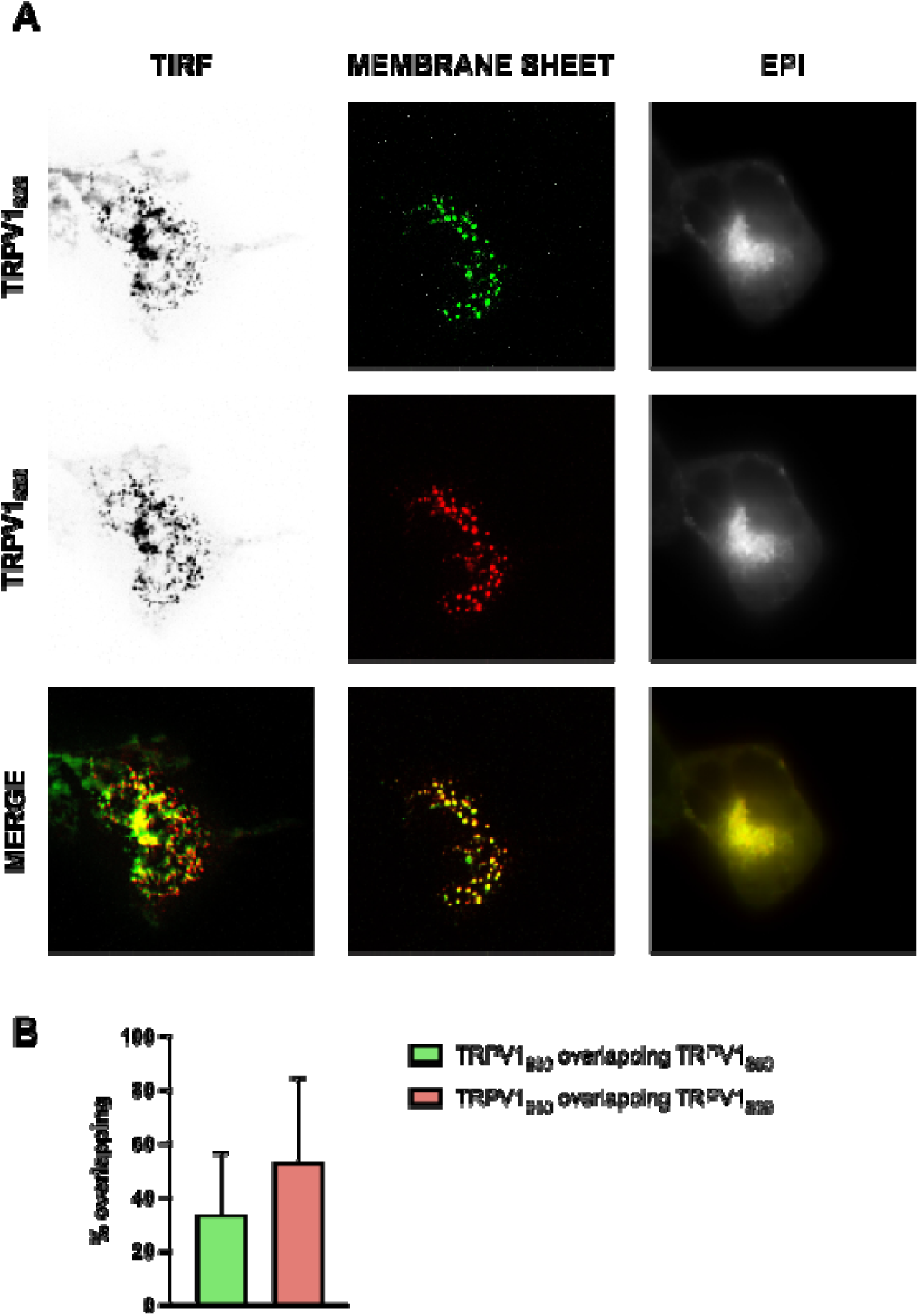
Total internal reflection fluorescence (TIRF) microscopy of cotransfected HEK293 cells with TRPV1_839_ and TRPV1_850_. **A.** Cells cotransfected with TRPV1_839_ and TRPV1_850_ show a puncta pattern at the TIRF plane that contrasts the diffuse pattern observed in EPI. The center panel displays the sharply defined puncta obtained in membrane sheets. **B.** Percentage of overlapping signal between TRPV1_839_ and TRPV1_850_.

### TRPV1_850_ modulates the cellular response to capsaicin

Imaging experiments suggest that the novel splice variant TRPV1_850_ affects the number of available TRPV1 channels at the plasma membrane. Thus, we assessed TRPV1_850_ functional characteristics using an electrophysiological approach (Figure 6A). Whole-cell patch-clamp recordings were performed first on HEK293 cells expressing TRPV1_839_ or TRPV1_850_ transcript variants (Figure 6B). Current versus voltage relationships showed modest outward rectifying currents of TRPV1_839_ channels in non-stimulated conditions and the expected robust capsaicin-evoked currents (Figure 6B-D). In contrast, the expression of TRPV1_850_ alone does not elicit currents (Figure 6B-D). This would agree with the reticular signal observed in TIRF microscopy experiments (Fig 4B and C).

**Figure 6.**
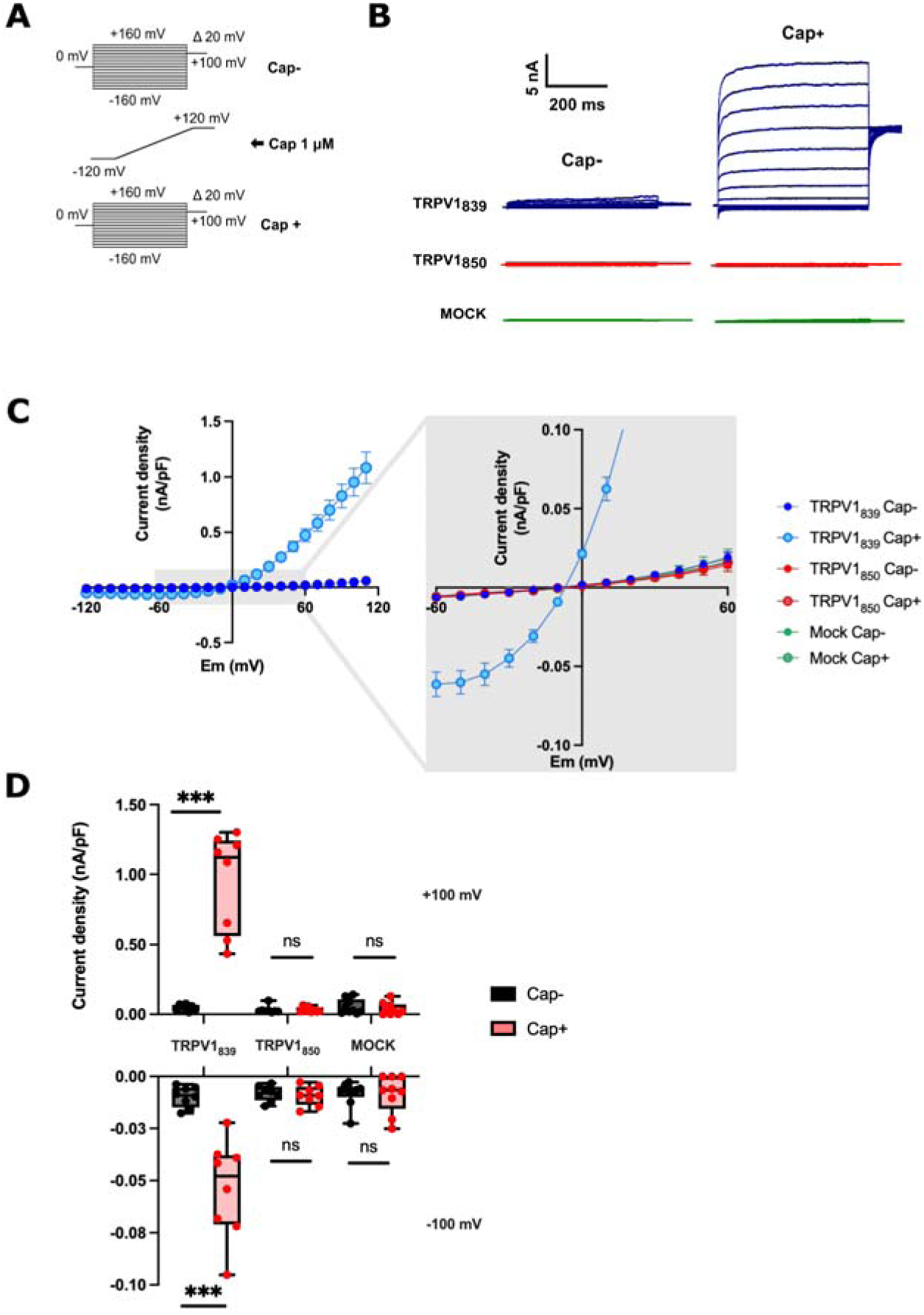
Whole-cell patch clamp experiments in transfected HEK293 cells with TRPV1_839_ or TRPV1_850_. **A.** Voltage step and ramp protocols. Capsaicin, 1 μM stimuli, were added after 1 minute during the ramp voltage protocol. **B.** Representative traces obtained from the voltage step protocol in the control condition and after the capsaicin 1 μM stimuli. **C.** Current-membrane potential relation in the control condition and after the capsaicin 1 μM stimuli. **D.** Current density at -100 and +100 mV in the control condition and after the capsaicin 1 μM stimuli. *** represents p value ≤ 0.001. MOCK represents an experiment in which cells were transfected with an empty vector.

On the other hand, the simultaneous expression of both splice variants results in outward rectifying currents similar to the ones obtained in transfected HEK293T cells with TRPV1_839_ alone (Fig. 7A-B). However, capsaicin stimulation (1uM) in the cotransfection condition renders significantly smaller currents (Fig. 7C). Despite the differences in the total current amplitude, the conductance versus voltage of both conditions, i.e., TRPV1_839_ alone or the cotransfection of both splice variants, is similar (Fig. 7D). These results support our findings from imaging experiments and, overall, suggest a putative role of TRPV1_850_ in controlling the number of channels rather than channel open probability and gating. Such a strategy might impact the cellular response to, for example, inflammatory nociceptive signals.

**Figure 7.**
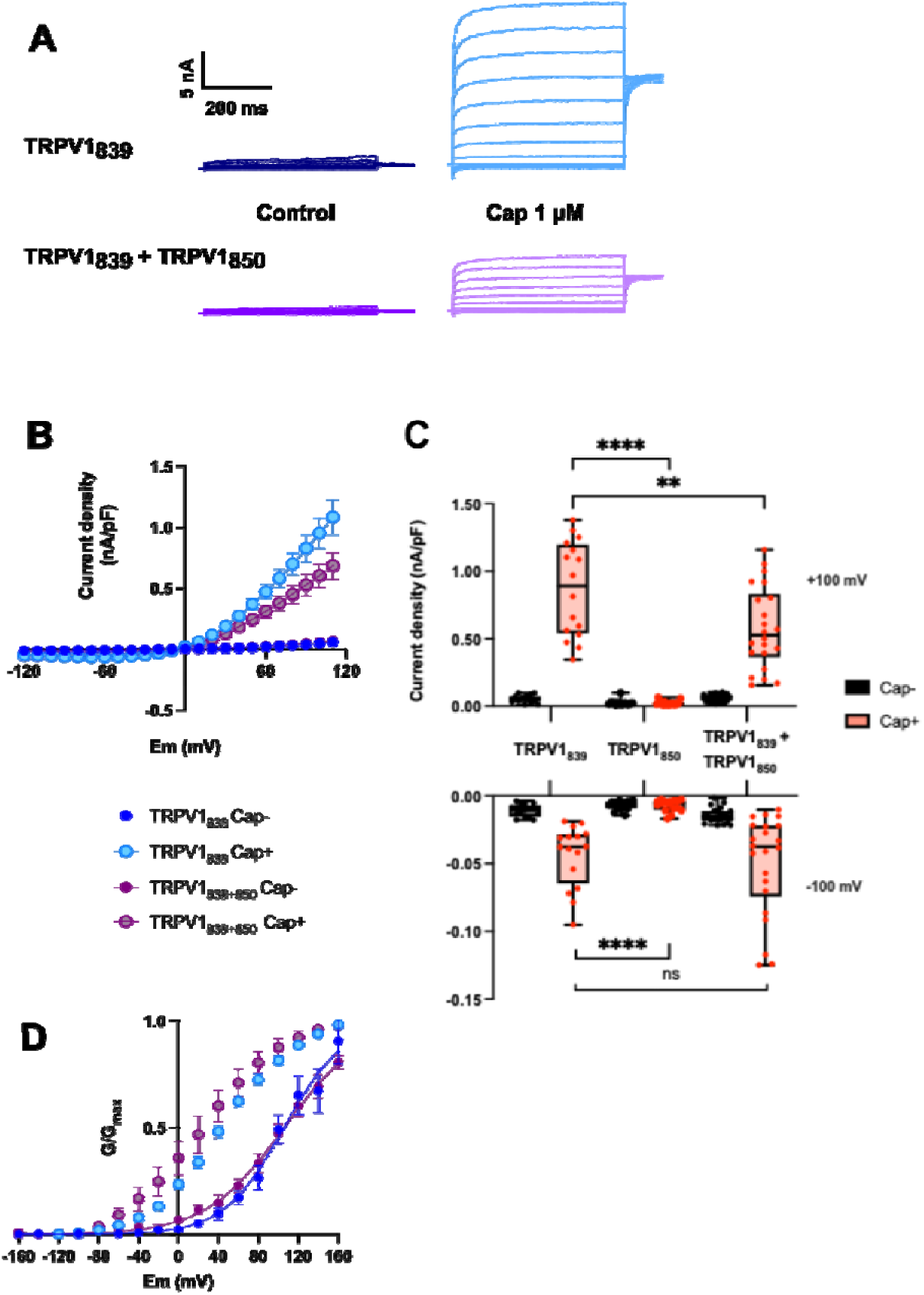
Whole-cell patch clamp experiments in cotransfected HEK293 cells with TRPV1_839_ and TRPV1_850_. **A.** Representative traces obtained from the voltage step protocol in the control condition and after the capsaicin 1 μM stimuli. **B.** Current-voltage relation in the control condition and after the capsaicin 1 μM stimuli. **C.** Current density at -100 and +100 mV in the control condition and after the capsaicin 1 μM stimuli. * represents a p-value ≤ 0.05. **D.** Normalized conductance (G/Gmax) versus voltage relationship. Experimental data were fitted to a Boltzmann function (see methods).

Splice variants or naturally occurring mutations in ion channels can significantly impact cellular physiology, often through dominant negative effects. A dominant negative effect occurs when a variant of a protein counteracts the function of the wild-type protein, often by forming non-functional or improperly functioning complexes. This phenomenon is particularly important in ion channels, which are typically multimeric complexes (Sanguinetti and Tristani-Firouzi 2006). The dominant negative effects of splice variants and natural mutants can arise through several mechanisms, such as impaired channel assembly, altered gating, mislocalization, or abnormal protein-protein interactions affecting channel regulatory strategies. Understanding the dominant negative effects of splice variants and natural mutants of ion channels is crucial for developing targeted therapies for various channelopathies.

Some examples of severe dominant negative effects include the HERG (human ether-à-go-go-related gene) potassium channel, which is critical for cardiac repolarization (Vandenberg et al. 2012). Some HERG splice variants can act as dominant negative mutants, impairing the function of the channel and, in turn, the cardiac muscle, leading to Long QT Syndrome, a condition predisposing individuals to arrhythmias and sudden cardiac death (London et al. 1997). Another example is the Cystic Fibrosis Transmembrane Conductance Regulator (CFTR), a chloride channel presenting numerous natural mutations that can be deleterious for channel function. One of the most common CFTR mutations is ΔF508, which can cause defective trafficking and function of the protein, leading to cystic fibrosis symptoms (Riordan 2008).

TRP channels are not the exception. For example, splice variants of the TRPM4 ion channel can act as dominant negatives, impairing the normal function of the channel and influencing cell signaling and volume regulation (Launay et al. 2002). Several splice variants have been reported for the TRPV1 ion channel (Schumacher and Eilers 2010). These splice variants introduce functional effects on the canonical channel (TRPV1_839_), modulating channel activity with either dominant negative or positive effects (Schumacher et al. 2000; Lu et al. 2005; Vos et al. 2006; Eilers et al. 2007; Tian et al. 2019).

The transient receptor potential vanilloid 1 (TRPV1) channel significantly mediates inflammatory responses by detecting harmful stimuli and initiating pain and inflammation pathways. While TRPV1 is integral to inflammatory responses across primate species, documented differences between catarrhine monkeys and other primates remain limited. However, variations in TRPV1 function and expression across species have been observed. For instance, certain species exhibit differences in TRPV1 sensitivity to temperature and chemical stimuli, which can influence inflammatory responses <u>(Shuba 2020)</u>. These functional disparities may be attributed to evolutionary adaptations to distinct environmental pressures and dietary habits. Further research is necessary to elucidate these potential differences and their physiological implications from a broader comparative biology perspective.

## Conclusions

Our study described an exonic sequence for the TRPV1 ion channel gene, which is only present in apes and Old World monkeys. From a structural perspective, incorporating this exon disrupts the sixth ankyrin domain in the N-terminal portion of the channel, which prevents this isoform from reaching the plasma membrane by itself. However, when this novel isoform (TRPV1_850_) is coexpressed with the canonical splice variant (TRPV1_839_), it does traffic to the plasma membrane. Experimental results show that smaller currents, compared to the condition when the canonical isoform is the only one expressed, are obtained in response to capsaicin stimulation when both isoforms are expressed. However, the conductance versus voltage of both conditions is similar on the slope and in their shift on the voltage axis after agonist stimulation. The intracellular location of the novel extra sequence, away from the selectivity filter, supports our underlying hypothesis, that the single channel conductance is unchanged between the homo and hetero multimers described here. Thus our results strongly suggest a putative regulatory mechanism by influencing the number of channels rather than channel mechanics of gating affecting open probability.

## Material and Methods

### DNA sequences

We manually annotated TRPV1 genes in representative species of all major groups of primates, i.e., apes, Old World monkeys, New World monkeys, tarsiers, and strepsirrhines (Supplementary Table S1). We also annotated TRPV1 channels in representative species of all other mammalian orders belonging to the superorder Euarchontoglires, i.e., scandentians, dermopterans, rodents, and lagomorphs (Supplementary Table S1). We first identified chromosomal fragments containing TRPV1 genes based on conserved synteny in the National Center for Biotechnology Information (NCBI) (Sayers et al. 2022). Once identified, chromosomal fragments, including the 5′ and 3′ flanking genes, were extracted. After extraction, we manually curated the existing annotation, or we annotate *de novo* by comparing human exon sequences (query sequence) to the species of which the genomic piece (subject sequence) is being analyzed using the program Blast2seq v2.5 with default parameters (Tatusova and Madden 1999).

### Identification of the splice variant incorporating the sixth exon, TRPV1_850_, at the transcriptional level

To identify the splice variant from the human TRPV1 gene (ENSG00000196689) that incorporates the sixth exon, we considered reads resulting from RNA-Seq experiments for which the origin of expression could be unequivocally associated with the microexon that characterizes the variant (ENST00000425167.6). For this, we selected Bioproject PRJEB4337 from the Sequence Read Archive SRA Database (Sayers et al. 2024), which has a broad sample of 27 healthy human tissues and 171 runs of RNA-Seq, with around 5 billion 101 bp paired-end reads, which had reported in the literature that TRPV1 channel gene is expressed (Sayers et al. 2024). We checked the read quality of the datasets using FastQC (https://www.bioinformatics.babraham.ac.uk/projects/fastqc/) and trimmed the reads using fastp v0.20.1 (Chen et al. 2018) with default parameters. This process trimmed the low-quality ends of the reads and sequencing adapters. We compared the original dataset qualities and the resulting datasets after trimming using MultiQC v1.9 (Ewels et al. 2016).

The next step is scanning all RNA-Seq data and filtering the reads of interest. We divided the scanning process into three steps to save resources and identify the subset of reads that align unambiguously over the microexon. In the first step, we mapped the reads to the sequence of the TRPV1 gene, filtering the majority of the reads. To this end, we downloaded the annotation of the TRPV1 gene transcript variants from Ensembl release v113 (ENSG00000196689 chr17:3565444-3609411; ENST00000425167) (Harrison et al. 2024). We used Ensembl annotation over NCBI’s because the annotation of the transcript variant of interest was not included in the GRCh38.p14 annotation files from NCBI’s Genome Database (GCF_000001405.40) (Sayers et al. 2024). The mapping was performed with hisat2 v.2.2.1 (Kim et al. 2019) with parameters -p 16 -a –-known-splicesites-infile splicesites.txt. We used the hisat2_extract_splice_sites.py script to extract known splice sites of the TRPV1 gene annotation and renumbered the start and end coordinates to relate the hisat2 results to the positions of the microexon. We used Samtools v1.11 (Danecek et al. 2021) to filter the reads in the region of the TRPV1 microexon (chr17:3589807-3588188 reverse strand).

In the second step, a visual inspection of the resulting filtered .sam file allowed us to manually select valid positive candidate reads aligning over the microexon. We visualized the reads in the region surrounding the microexon (chr17:3589807-3588188 reverse strand) using Samtools tview and the Integrated Genome Viewer, IGV v2.18.2 (Robinson et al. 2011). The criteria for selection were uniquely mapped reads spliced on one or both exons surrounding the sixth exon. The coordinates of read splicing had to match the exon annotation coordinates.

In the third step we mapped the selected reads over the whole human genome (GRCh38.p14) without using the known splice sites feature, eliminating remaining false positives and ensuring that the reads aligned uniquely and were spliced among the surrounding exons. For this, the human genome (GRCh38.p14) sequence was downloaded from NCBI FTP (https://ftp.ncbi.nlm.nih.gov/genomes/all/GCF/000/001/405/GCF_000001405.40_GRCh38.p14/GCF_000001405.40_GRCh38.p14_genomic.fna.gz). The selected reads were mapped using hisat2 v.2.2.1. The mapping quality (MAPQ) and CIGAR of the reads in the .sam file were inspected to confirm that the reads originated from the region to which they aligned.

### cDNA constructs

Full-length hTRPV1 coding sequences were cloned in pEGFP-C1 for heterologous expression in mammalian cells. DNA constructs replacing EGFP by HaloTag (Los et al. 2008) were synthesized by Genescript. Halo-Sec61b was kindly provided by Jennifer Lippincott-Schwartz (HHMI, Janelia).

### Cell culture and expression

HEK293T cells (ATCC) were cultured in DMEM and supplied with 10% FBS (both Invitrogen) and 1% PenStrep (Invitrogen). Cells were plated on poly-L-lysine (Sigma-Aldrich) coated coverslips and transfected using poly-jet (Signagen) according to the manufacturer’s instructions. Both imaging and electrophysiology recordings were performed 24–48 hours after transfection. Low transfection was accomplished at 12–18 h when required.

### Solutions

Unless stated otherwise, the pipette solution contained (in mM): 105 CsF, 35 NaCl, 10 EGTA, 8 KCl, 0.1 CaCl_2_ and 10 HEPES, pH 7.4. The bath solution contained (in mM): 140 NaCl, 2 CaCl_2_, 5 KCl, 2 MgCl_2_, 10 HEPES, and 8 glucose, pH 7.4. Imaging was performed in a solution containing (in mM): 140 NaCl, 2 CaCl_2_, 5 KCl, 2 MgCl_2_, 10 HEPES, and 8 glucose, pH 7.4.

### Electrophysiology

Whole-cell currents were obtained from transiently transfected HEK293T cells. Gigaseals were formed using 2- to 4-MΩ borosilicate pipettes (A-M Systems). Junction potentials (6.8 mV) were corrected for CsF/NaCl solutions. Seal resistance was >0.7 GΩ in all cases. Series resistance and cell capacitance were analogically compensated directly on the amplifier. Whole-cell voltage clamp was performed, and the macroscopic currents were acquired at 10 kHz and filtered at 2 kHz. Data was acquired using an Axopatch 200B (Molecular Devices) or a Model 2400 patch clamp amplifier (A-M Systems). Signals were digitized on a 16-bit Digidata 1322A (Molecular Devices).

Current-voltage relationships were studied using a voltage step protocol of 400 ms each step from -160 mV to +160 mV using 20 mV intervals, followed by a 400-ms linear ramp from -120 to +120 mV. The time interval between each ramp was 2 s, and 1 μM capsaicin was applied during the ramp protocol, followed by another voltage step protocol in the presence of capsaicin (see scheme on Figure 6A). Currents are presented in terms of densities (pA/pF). The conductance versus voltage relations were fitted to a Boltzmann distribution of the form G = Gmax/(1 + exp - [zF(V-V0.5)/RT]), where z is the voltage dependency, V0.5 is the half activation voltage, and Gmax is the maximum conductance.

### Total internal reflection fluorescence (TIRF) microscopy

HEK293T cells were imaged using a through-the-objective TIRF microscope fed by 488 nm and 561 nm DPSS laser lines (Coherent) built on a Nikon TiS. Both lasers were focused onto a 10 µm optical fiber and transmitted via the rear illumination port. Digitally synchronized mechanical shutters (Vincent Associates) controlled exposure times through a high-numerical aperture objective (60x, 1.49 NA, oil; Nikon). Fluorescence emission was recorded by a CMOS detector (BFY-U3-51S5M, FLIR). Imaging was performed at 2 Hz using a micro-manager (Universal Imaging).

### Membrane sheet preparation

Untransfected HEK293 cells were washed twice in an ice-cold imaging buffer, before being subjected to 3 sonication pulses of 1s each at 40% power (Cole Palmer 4710 Ultrasonics Homogenizer) directly on coverslips. Cells were washed twice in the imaging buffer prior imaging.

### Image analysis

Images were analyzed using Image J (NIH), and pre-treatment consisted of an average of 20 frames and background subtraction. The WEKA Segmentation plug-in (Arganda-Carreras et al., 2017) was trained and used to detect signal spots at the membrane (i.e., puncta, and clusters). Manders coefficients were obtained using the JaCoP Colocalization Analysis plug-in (Bolte and Cordeliéres, 2006) for co-transfected cell images with default parameters.

## Acknowledgments

This work was supported by the Fondo Nacional de Desarrollo Científico y Tecnológico from Chile, FONDECYT 1210471 to JCO, and 1241753 to SEB. The Genotype-Tissue Expression (GTEx) Project was supported by the Common Fund of the Office of the Director of the National Institutes of Health and by NCI, NHGRI, NHLBI, NIDA, NIMH, and NINDS. The data used for the analyses described in this manuscript were obtained from the Adult GTEx Portal on 10/22/2024.

## Author contributions

All authors contributed to the work presented in this paper. SEB and JCO designed research; SC, DCB, EM, JY, KZ, CM, SBV, RD, and GR performed research; SC, DCB, EM, JY, KZ, CM, SBV, RD, and GR analyzed data; and SEB and JCO wrote the paper. GR, RM and JY revised versions of the manuscript.

## Data Availability

Data and supplementary material are available online at https://github.com/opazolab/TRPV1

## Literature

Alekseyenko AV, Kim N, Lee CJ. 2007. Global analysis of exon creation versus loss and the role of alternative splicing in 17 vertebrate genomes. RNA 13:661–670.

Berget SM, Moore C, Sharp PA. 1977. Spliced segments at the 5’ terminus of adenovirus 2 late mRNA. Proc. Natl. Acad. Sci. U. S. A. 74:3171–3175.

Brauchi SE. 2023. Cold blooded vertebrates help unveil a heat-dependent trigger. Trends Neurosci. 46:781–782.

Caterina MJ, Schumacher MA, Tominaga M, Rosen TA, Levine JD, Julius D. 1997. The capsaicin receptor: a heat-activated ion channel in the pain pathway. Nature 389:816–824.

Chakrabarty B, Parekh N. 2022. Sequence and Structure-Based Analyses of Human Ankyrin Repeats. Molecules [Internet] 27. Available from: 10.3390/molecules27020423

Chen S, Zhou Y, Chen Y, Gu J. 2018. fastp: an ultra-fast all-in-one FASTQ preprocessor. Bioinformatics 34:i884–i890.

Chow LT, Gelinas RE, Broker TR, Roberts RJ. 1977. An amazing sequence arrangement at the 5’ ends of adenovirus 2 messenger RNA. Cell 12:1–8.

Danecek P, Bonfield JK, Liddle J, Marshall J, Ohan V, Pollard MO, Whitwham A, Keane T, McCarthy SA, Davies RM, et al. 2021. Twelve years of SAMtools and BCFtools. Gigascience [Internet] 10. Available from: 10.1093/gigascience/giab008

Desiere F, Deutsch EW, King NL, Nesvizhskii AI, Mallick P, Eng J, Chen S, Eddes J, Loevenich SN, Aebersold R. 2006. The PeptideAtlas project. Nucleic Acids Res 34:D655–D658.

Eilers H, Lee S-Y, Hau CW, Logvinova A, Schumacher MA. 2007. The rat vanilloid receptor splice variant VR.5’sv blocks TRPV1 activation. Neuroreport 18:969–973.

Ewels P, Magnusson M, Lundin S, Käller M. 2016. MultiQC: summarize analysis results for multiple tools and samples in a single report. Bioinformatics 32:3047–3048.

Fagerberg L, Hallström BM, Oksvold P, Kampf C, Djureinovic D, Odeberg J, Habuka M, Tahmasebpoor S, Danielsson A, Edlund K, et al. 2014. Analysis of the human tissue-specific expression by genome-wide integration of transcriptomics and antibody-based proteomics. Mol Cell Proteomics 13:397–406.

Ferrandiz-Huertas C, Mathivanan S, Wolf CJ, Devesa I, Ferrer-Montiel A. 2014. Trafficking of ThermoTRP Channels. Membranes (Basel) 4:525–564.

Gaudet R. 2008. A primer on ankyrin repeat function in TRP channels and beyond. Mol. Biosyst. 4:372–379.

Guan W, Orellana KG, Stephens RF, Zhorov BS, Spafford JD. 2022. A lysine residue from an extracellular turret switches the ion preference in a Cav3 T-Type channel from calcium to sodium ions. J. Biol. Chem. 298:102621.

Harrison PW, Amode MR, Austine-Orimoloye O, Azov AG, Barba M, Barnes I, Becker A, Bennett R, Berry A, Bhai J, et al. 2024. Ensembl 2024. Nucleic Acids Res 52:D891–D899.

Hori S, Tateyama M, Shirai T, Kubo Y, Saitoh O. 2023. Two single-point mutations in Ankyrin Repeat one drastically change the threshold temperature of TRPV1. Nat. Commun. 14:2415.

Katoh K, Rozewicki J, Yamada KD. 2019. MAFFT online service: multiple sequence alignment, interactive sequence choice and visualization. Brief. Bioinform. 20:1160–1166.

Kim D, Paggi JM, Park C, Bennett C, Salzberg SL. 2019. Graph-based genome alignment and genotyping with HISAT2 and HISAT-genotype. Nat Biotechnol 37:907–915.

Kondrashov FA, Koonin EV. 2001. Origin of alternative splicing by tandem exon duplication. Hum. Mol. Genet. 10:2661–2669.

Kopelman NM, Lancet D, Yanai I. 2005. Alternative splicing and gene duplication are inversely correlated evolutionary mechanisms. Nat. Genet. 37:588–589.

Kumar S, Suleski M, Craig JM, Kasprowicz AE, Sanderford M, Li M, Stecher G, Hedges SB. 2022. TimeTree 5: An Expanded Resource for Species Divergence Times. Mol. Biol. Evol. [Internet] 39. Available from: 10.1093/molbev/msac174

Ladrón-de-Guevara E, Dominguez L, Rangel-Yescas GE, Fernández-Velasco DA, Torres-Larios A, Rosenbaum T, Islas LD. 2020. The Contribution of the Ankyrin Repeat Domain of TRPV1 as a Thermal Module. Biophys. J. 118:836–845.

Launay P, Fleig A, Perraud AL, Scharenberg AM, Penner R, Kinet JP. 2002. TRPM4 is a Ca2+-activated nonselective cation channel mediating cell membrane depolarization. Cell 109:397–407.

Liang L, Fazel Darbandi S, Pochareddy S, Gulden FO, Gilson MC, Sheppard BK, Sahagun A, An J-Y, Werling DM, Rubenstein JLR, et al. 2021. Developmental dynamics of voltage-gated sodium channel isoform expression in the human and mouse brain. Genome Med. 13:135.

Lishko PV, Procko E, Jin X, Phelps CB, Gaudet R. 2007. The ankyrin repeats of TRPV1 bind multiple ligands and modulate channel sensitivity. Neuron 54:905–918.

Liu Z, Zhu C, Steinmetz LM, Wei W. 2023. Identification and quantification of small exon-containing isoforms in long-read RNA sequencing data. Nucleic Acids Res 51:e104.

London B, Trudeau MC, Newton KP, Beyer AK, Copeland NG, Gilbert DJ, Jenkins NA, Satler CA, Robertson GA. 1997. Two isoforms of the mouse ether-a-go-go-related gene coassemble to form channels with properties similar to the rapidly activating component of the cardiac delayed rectifier K+ current. Circ Res 81:870–878.

Los GV, Encell LP, McDougall MG, Hartzell DD, Karassina N, Zimprich C, Wood MG, Learish R, Ohana RF, Urh M, et al. 2008. HaloTag: a novel protein labeling technology for cell imaging and protein analysis. ACS Chem Biol 3:373–382.

Lu G, Henderson D, Liu L, Reinhart PH, Simon SA. 2005. TRPV1b, a functional human vanilloid receptor splice variant. Mol Pharmacol 67:1119–1127.

Mason VC, Li G, Minx P, Schmitz J, Churakov G, Doronina L, Melin AD, Dominy NJ, Lim NT-L, Springer MS, et al. 2016. Genomic analysis reveals hidden biodiversity within colugos, the sister group to primates. Sci Adv 2:e1600633.

Merkin JJ, Chen P, Alexis MS, Hautaniemi SK, Burge CB. 2015. Origins and impacts of new mammalian exons. Cell Rep. 10:1992–2005.

Mosavi LK, Cammett TJ, Desrosiers DC, Peng Z-Y. 2004. The ankyrin repeat as molecular architecture for protein recognition. Protein Sci. 13:1435–1448.

Ozaita A, Martone ME, Ellisman MH, Rudy B. 2002. Differential subcellular localization of the two alternatively spliced isoforms of the Kv3.1 potassium channel subunit in brain. J. Neurophysiol. 88:394–408.

Parada GE, Munita R, Georgakopoulos-Soares I, Fernandes HJR, Kedlian VR, Metzakopian E, Andres ME, Miska EA, Hemberg M. 2021. MicroExonator enables systematic discovery and quantification of microexons across mouse embryonic development. Genome Biol 22:43.

Perelman P, Johnson WE, Roos C, Seuánez HN, Horvath JE, Moreira MAM, Kessing B, Pontius J, Roelke M, Rumpler Y, et al. 2011. A molecular phylogeny of living primates. PLoS Genet. 7:e1001342.

Poteser M, Leitinger G, Pritz E, Platzer D, Frischauf I, Romanin C, Groschner K. 2016. Live-cell imaging of ER-PM contact architecture by a novel TIRFM approach reveals extension of junctions in response to store-operated Ca-entry. Sci. Rep. 6:35656.

Riordan JR. 2008. CFTR function and prospects for therapy. Annu. Rev. Biochem. 77:701–726.

Rivera B, Orellana-Serradell O, Servili E, Santos R, Brauchi S, Cerda O. 2024. The odyssey of the TR(i)P journey to the cellular membrane. Front Cell Dev Biol 12:1414935.

Robinson JT, Thorvaldsdóttir H, Winckler W, Guttman M, Lander ES, Getz G, Mesirov JP. 2011. Integrative genomics viewer. Nat Biotechnol 29:24–26.

Rosenbaum T, Islas LD. 2023. Molecular Physiology of TRPV Channels: Controversies and Future Challenges. Annu Rev Physiol 85:293–316.

Sanguinetti MC, Tristani-Firouzi M. 2006. hERG potassium channels and cardiac arrhythmia. Nature 440:463–469.

Sayers EW, Beck J, Bolton EE, Brister JR, Chan J, Connor R, Feldgarden M, Fine AM, Funk K, Hoffman J, et al. 2024. Database resources of the National Center for Biotechnology Information in 2025. Nucleic Acids Res [Internet]. Available from: 10.1093/nar/gkae979

Sayers EW, Bolton EE, Brister JR, Canese K, Chan J, Comeau DC, Connor R, Funk K, Kelly C, Kim S, et al. 2022. Database resources of the national center for biotechnology information. Nucleic Acids Res. 50:D20–D26.

Schumacher MA, Eilers H. 2010. TRPV1 splice variants: structure and function. Front Biosci (Landmark Ed) 15:872–882.

Schumacher MA, Moff I, Sudanagunta SP, Levine JD. 2000. Molecular cloning of an N-terminal splice variant of the capsaicin receptor. Loss of N-terminal domain suggests functional divergence among capsaicin receptor subtypes. J Biol Chem 275:2756–2762.

Senatore A, Guan W, Boone AN, Spafford JD. 2014. T-type channels become highly permeable to sodium ions using an alternative extracellular turret region (S5-P) outside the selectivity filter. J. Biol. Chem. 289:11952–11969.

Senning EN, Gordon SE. 2015. Activity and Ca^2+^ regulate the mobility of TRPV1 channels in the plasma membrane of sensory neurons. Elife 4:e03819.

Shao Y, Zhou L, Li F, Zhao L, Zhang B-L, Shao F, Chen J-W, Chen C-Y, Bi X, Zhuang X-L, et al. 2023. Phylogenomic analyses provide insights into primate evolution. Science 380:913–924.

Sharif Naeini R, Witty M-F, Séguéla P, Bourque CW. 2006. An N-terminal variant of Trpv1 channel is required for osmosensory transduction. Nat Neurosci 9:93–98.

Singh P, Ahi EP. 2022. The importance of alternative splicing in adaptive evolution. Mol. Ecol. 31:1928–1938.

Szallasi A, Cortright DN, Blum CA, Eid SR. 2007. The vanilloid receptor TRPV1: 10 years from channel cloning to antagonist proof-of-concept. Nat Rev Drug Discov 6:357–372.

Tatusova TA, Madden TL. 1999. BLAST 2 Sequences, a new tool for comparing protein and nucleotide sequences. FEMS Microbiol. Lett. 174:247–250.

Tian Q, Hu J, Xie C, Mei K, Pham C, Mo X, Hepp R, Soares S, Nothias F, Wang Y, et al. 2019. Recovery from tachyphylaxis of TRPV1 coincides with recycling to the surface membrane. Proc Natl Acad Sci U S A 116:5170–5175.

Toro CA, Eger S, Veliz L, Sotelo-Hitschfeld P, Cabezas D, Castro MA, Zimmermann K, Brauchi S. 2015. Agonist-dependent modulation of cell surface expression of the cold receptor TRPM8. J Neurosci 35:571–582.

UniProt Consortium. 2023. UniProt: the Universal Protein Knowledgebase in 2023. Nucleic Acids Res. 51:D523–D531.

Vandenberg JI, Perry MD, Perrin MJ, Mann SA, Ke Y, Hill AP. 2012. hERG K(+) channels: structure, function, and clinical significance. Physiol Rev 92:1393–1478.

Vázquez E, Valverde MA. 2006. A review of TRP channels splicing. Semin Cell Dev Biol 17:607–617.

Verta J-P, Jacobs A. 2022. The role of alternative splicing in adaptation and evolution. Trends Ecol. Evol. 37:299–308.

Vos MH, Neelands TR, McDonald HA, Choi W, Kroeger PE, Puttfarcken PS, Faltynek CR, Moreland RB, Han P. 2006. TRPV1b overexpression negatively regulates TRPV1 responsiveness to capsaicin, heat and low pH in HEK293 cells. J Neurochem 99:1088–1102.

Wang C, Hu H-Z, Colton CK, Wood JD, Zhu MX. 2004. An alternative splicing product of the murine trpv1 gene dominant negatively modulates the activity of TRPV1 channels. J Biol Chem 279:37423–37430.

Yu H, Li M, Sandhu J, Sun G, Schnable JC, Walia H, Xie W, Yu B, Mower JP, Zhang C. 2022. Pervasive misannotation of microexons that are evolutionarily conserved and crucial for gene function in plants. Nat Commun 13:820.

Ferrandiz-Huertas C, Mathivanan S, Wolf CJ, Devesa I, Ferrer-Montiel A. Trafficking of ThermoTRP Channels. Membranes. 2014; 4(3):525–564.

Tian Q, Hu J, Xie C, Mei K, Pham C, Mo X, Hepp R, Soares S, Nothias F, Wang Y, Liu Q, Cai F, Zhong B, Li D, Yao J. Recovery from tachyphylaxis of TRPV1 coincides with recycling to the surface membrane. Proc. Natl. Acad. Sci. U.S.A. 116(11):5170–5175.

